# Passing the message: representation transfer in modular balanced networks

**DOI:** 10.1101/716118

**Authors:** Barna Zajzon, Sepehr Mahmoudian, Abigail Morrison, Renato Duarte

## Abstract

Neurobiological systems rely on hierarchical and modular architectures to carry out intricate computations using minimal resources. A prerequisite for such systems to operate adequately is the capability to reliably and efficiently transfer information across multiple modules. Here, we study the features enabling a robust transfer of stimulus representations in modular networks of spiking neurons, tuned to operate in a balanced regime. To capitalize on the complex, transient dynamics that such networks exhibit during active processing, we apply reservoir computing principles and probe the systems’ computational efficacy with specific tasks. Focusing on the comparison of random feed-forward connectivity and biologically inspired topographic maps, we find that, in a sequential set-up, structured projections between the modules are strictly necessary for information to propagate accurately to deeper modules. Such mappings not only improve computational performance and efficiency, they also reduce response variability, increase robustness against interference effects, and boost memory capacity. We further investigate how information from two separate input streams is integrated and demonstrate that it is more advantageous to perform non-linear computations on the input locally, within a given module, and subsequently transfer the result downstream, rather than transferring intermediate information and performing the computation downstream. Depending on how information is integrated early on in the system, the networks achieve similar task-performance using different strategies, indicating that the dimensionality of the neural responses does not necessarily correlate with nonlinear integration, as predicted by previous studies. These findings highlight a key role of topographic maps in supporting fast, robust and accurate neural communication over longer distances. Given the prevalence of such structural feature, particularly in the sensory systems, elucidating their functional purpose remains an important challenge towards which this work provides relevant, new insights. At the same time, these results shed new light on important requirements for designing functional hierarchical spiking networks.

**Author summary:** To interact with the external environment in real-time, cortical microcircuits must employ efficient and reliable mechanisms for passing information between different modules and for integrating input from multiple sources. In this study we investigate, from a functional perspective, how structural features influence these mechanisms in the context of stimulus representation, integration and transfer in modular spiking networks. We demonstrate that biologically plausible patterned connectivity, inspired by cortical topographic maps, is necessary for information to propagate across multiple modules in a useful manner. Compared to purely random projections, topographic maps improve computational performance and efficiency considerably, leading to more stable responses and increased robustness against interference effects. In addition, architectural specificities also play an important role in how networks combine information from different sources. Our results suggest that early and local integration of input streams enables more accurate non-linear computations in deeper modules than first transferring intermediate representations and only then performing the computations downstream. These findings demonstrate that the wiring architecture can profoundly impact information transmission and integration in neuronal circuits, with structured connectivity in form spatially segregated topographic projections playing a potentially key role in sustaining accurate cortical communication over multiple networks.

## Introduction

Cortical information processing relies on a distributed functional architecture comprising multiple, specialized modules arranged in complex, but stereotyped networks (see, e.g. [1–3]. Structural organizational principles are noticeable at different scales and impose strong constraints on the systems’ functionality [4], while simultaneously suggest a certain degree of uniformity and a close relation between structure and function [5, 6].

On the lower levels of cortical processing, peripheral signals conveying sensory information need to be adequately routed, their content represented and integrated with internal, ongoing processes [7] (based on both local and long-range interactions) as well as non-sensory signals such as attention [8], expectation [9] or reward [10]. A prerequisite for processing across such large distributed systems is therefore the ability to suitably represent relevant stimulus features, and transfer these representations in a reliable and efficient manner through various processing modules. Additionally, cortical areas are arranged in a functional hierarchy [3, 11, 12], whereby *higher*, more anterior, regions show sensitivity to increasingly complex and abstract features.

The computational benefits of such hierarchical feature aggregation and modular specialization have been consistently demonstrated in the domain of artificial neural networks [13], with a primary focus on spatial and/or spectral features. However, given that, to a first approximation, cortical systems are recurrent networks of spiking neurons, temporal dynamics and the ability to continuously represent and process spatio-temporal information are fundamental aspects of neural computation. The majority of previous studies on spatio-temporal processing with spiking neural networks have either focused on local information processing without considering the role of, or mechanisms for, modular specialization (e.g. [14]), or on the properties of signal transmission within one or across multiple neuronal populations regardless of their functional context ([15–20], but see, e.g. [21, 22] for counter-examples).

In order to quantify transmission accuracy and, implicitly, information content, these studies generally look either at the stable propagation of synchronous spiking activity [17] or asynchronous firing rates [18]. The former involves the temporally precise transmission of pulse packets (or spike volleys) aided by increasingly synchronous responses in multi-layered feed-forward networks (so-called “synfire chains”); the later refers to the propagation of asynchronous activity and assumes that information is contained and forwarded in the fidelity of the firing rates of individual neurons or certain sub-populations. An alternative approach was recently taken by [20], in which signal propagation was analyzed in a large-scale cortical model and elevated firing rates across areas were considered a signature of successful information transmission. However, no transformations on the input signals were carried out. Thus, a systematic analysis that considers both computation within a module and the transmission of computational results to downstream modules remains to be established.

In this study, we hypothesize that biophysically-based architectural features (modularity and topography) impose critical functional constraints on the reliability of information transmission, aggregation and processing. To address some of the issues and limitations highlighted above, we consider a system composed of multiple interconnected modules, each of which is realized as a recurrently coupled network of spiking neurons, acting as a state-dependent processing reservoir whose high-dimensional transient dynamics supports online computation with fading memory, allowing simple readouts such as linear classifiers to learn a large set of input–output relations [23]. Through the effect of the nonlinear nodes and their recurrent interactions, each module projects its inputs to a high dimensional feature space retaining time course information in the transient network responses. By connecting such spiking neural network modules, we uncover the architectural constraints necessary to enable a reliable transfer of stimulus representations from one module to the next. Using such a *reservoir computing* (RC) approach [24], the transmitted signals are conferred functional meaning and the circuits’ information processing capabilities can be probed in various computational contexts. Preliminary results from this approach have been presented in a conference proceedings [25].

Our results demonstrate that the connectivity structure between the modules strongly affects the transmission efficacy. We contrast random projections with biologically-inspired topographic maps, which are particularly prominent in sensory systems [26] and have been associated with a variety of important functional roles, ranging from information segregation and transmission along sensory pathways [27, 28] to spatio-temporal feature aggregation [29]. Additionally, conserved topography was shown to support the development of stable one-to-one mappings between abstract cognitive representations in higher cortical regions [30]. We show that incorporating such structured projections between the modules facilitates the reliable transmission of information, improving the overall computational performance. Such ordered mappings lead to lower-dimensional neural responses, allowing a more stable and efficient propagation of the input throughout the modules while enabling a computationally favorable dynamic regime. These results suggest that, while random connectivity can be applied for local processing within a module or between a few populations, accurate and robust information transmission over longer distances benefits from spatially segregated pathways and thus offers a potential functional interpretation for the existence of conserved topographic maps patterning cortical mesoscopic connectivity.

## Methods

### Network architecture

We model systems composed of multiple sub-networks (modules). Each module is a balanced random network (see, e.g. [31]), i.e., a sparsely and randomly connected recurrent network containing *N* = 10000 leaky integrate-and-fire neurons (described below), sub-divided into *N*^E^ = 0.8*N* excitatory and *N*^I^ = 0.2*N* inhibitory populations. Neurons make random recurrent connections within a module with a fixed probability common for all modules, *ϵ* = 0.1, such that on average each neuron in every module receives recurrent input from *K*_E_ = *ϵN*^E^ excitatory and *K*_I_ = *ϵN*^I^ inhibitory local synapses.

For simplicity, all projections between the modules are considered to be purely feed-forward and excitatory. Specifically, population *E_i_* in module *M_i_* connects, with probability *p*_ff_, to both populations *E*_*i*+1_ and *I*_*i*+1_ in subsequent module *M*_*i*+1_. This way, every neuron in *M*_*i*+1_ receives an additional source of excitatory input, mediated via *K*_*M*_*i*+1__ = *p*_ff_*N*^E^ synapses (see Fig 1).

**Fig 1.**
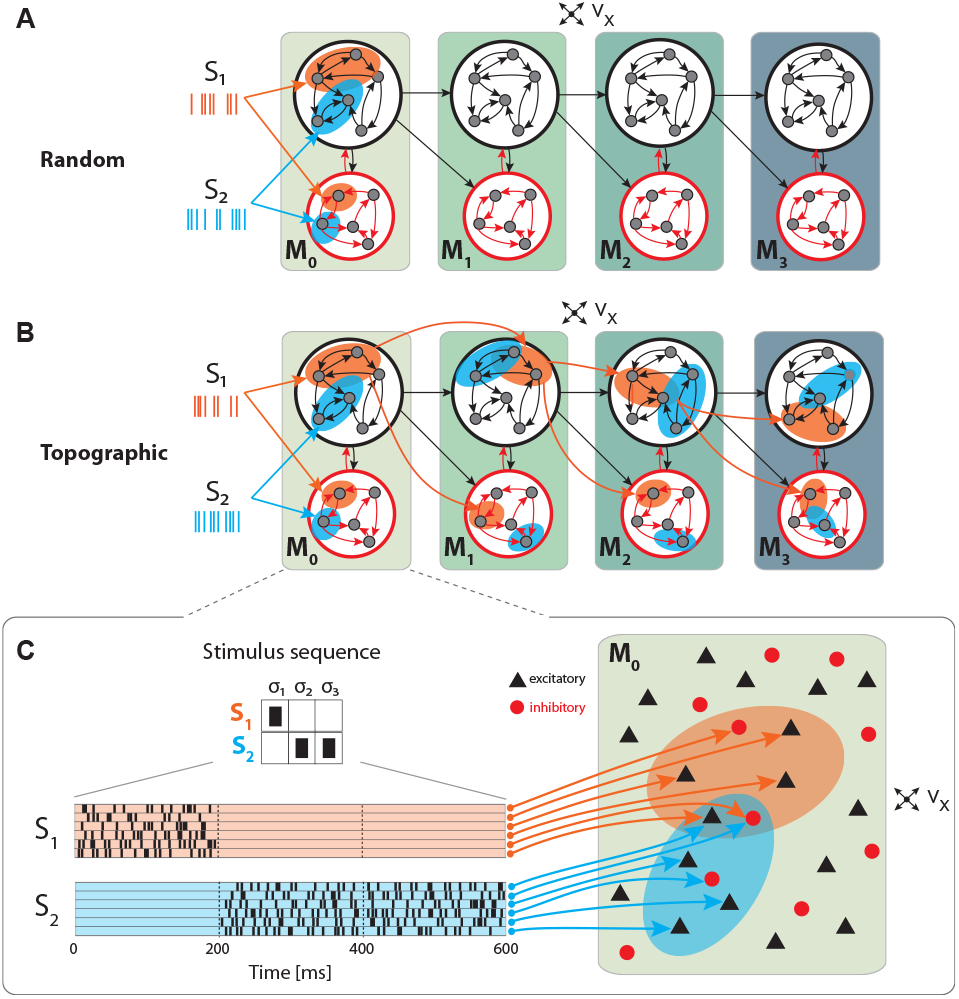
Schematic overview of the sequential setup and input stimuli. Networks are composed of four modules with identical internal structure, with random (**A**) or topographically structured (**B**) feed-forward projections. Structured stimuli drive specific, randomly selected sub-populations in *M*_0_. For stimulus *S*_1_, the topographic projections (**B**, orange arrows) between the modules are represented explicitly in addition to the corresponding stimulus-specific sub-populations (orange ellipses), whereas for *S*_2_ only the sub-populations are depicted (blue ellipses). The black feed-forward arrows depict the remaining sparse random connections from neurons that are not part of any stimulus-specific cluster. **C**: Illustrative example of the input encoding scheme: a symbolic input sequence of length *T* (3 in this example), containing |*S*| different, randomly ordered stimuli (*S* = {*S*_1_, *S*_2_}), is encoded into a binary matrix of dimensions |*S*| × *T*. Each stimulus is then converted into a set of 800 Poissonian spike trains of fixed duration (200 ms) and rate *ν*_stim_ and delivered to a subset of *ϵN*^E^ excitatory and *ϵN*^I^ inhibitory neurons.

To place the system in a responsive regime, all neurons in each module further receive stochastic external input (background noise) from *K*_x_ = *p*_x_*N*^x^ synapses. We set *N*^x^ = *N*^E^, as it is commonly assumed that the number of background input synapses modeling local and distant cortical input is in the same range as the number of recurrent excitatory connections [15, 31, 32].

In order to preserve the operating point of the different sub-networks, we scale the total input from sources external to each module to ensure that all neurons (regardless of their position in the setup) receive, on average, the same amount of excitatory drive. Whereas *p*_x_ = *ϵ* holds in the first (input) module, *M*_0_, the connection densities for deeper modules are chosen such that *p*_ff_ + *p*_x_ = *ϵ*, with *p*_ff_ = 0.75*ϵ* and *p*_x_ = 0.25*ϵ*, yielding a ratio of 3:1 between the number of feed-forward and background synapses.

For a complete, tabular description of the models and model parameters used throughout this study, see Supplementary Tables S1 and S2.

### Structured feed-forward connectivity

We explore the functional role of long-range connectivity profiles by investigating and comparing networks with random (Fig 1A) and topographically structured feed-forward projections (Fig 1B).

To build systems with topographic projections in a principled, but simple, manner, a network with random recurrent and feed-forward connectivity (as described in the previous section) is modified by systematically assigning sub-groups of stimulus-specific neurons in each module. Each of these then connects only to the corresponding sub-group across the different modules. More specifically, each stimulus *S_k_* projects onto a randomly chosen subset of 800 excitatory and 200 inhibitory neurons in *M*_0_ (input module), denoted 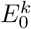 and 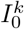. The connections from 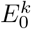 to module *M*_1_ are then rewired such that neurons in 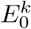 project, with probability *p*_ff_, exclusively to similarly chosen stimulus-specific neurons 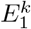 and 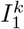. These sub-populations in *M*_1_ thus extend the topographic map associated with stimulus *S_k_*. By repeating these steps throughout the system, we ensure that each stimulus is propagated through a specific pathway while inter-module projections from neurons not belonging to any topographic map remain unchanged (random). This connectivity scheme is illustrated for stimulus *S*_1_ in Fig 1B.

It is worth noting that, as the stimulus-specific sub-populations are randomly chosen, overlaps occur (depending on the total number of stimuli). By allowing multiple feed-forward synaptic connections between neurons that are part of different clusters, the effective connection density along the topographic maps (*p*_ff_) is slightly increased compared with the random case (from 0.075 to 0.081). Any given neuron belongs to at most three different maps, ensuring that information transmission is not heavily biased by only a few strong connections. The average overlap between maps, measured as the mean fraction of neurons shared between any two maps, was 0.61. These values are representative for all sequential setups, unless stated otherwise.

### Neuron and synapse model

The networks are composed of leaky integrate-and-fire (LIF) neurons, with fixed voltage threshold and conductance-based, static synapses. The dynamics of the membrane potential *V_i_* for neuron *i* follows:

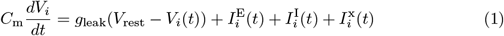

where the leak-conductance is given by *g*_leak_, and 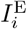 and 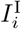 represent the total excitatory and inhibitory synaptic input currents, respectively. We assume the external background input, denoted by 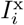, to be excitatory (all parameters equal to recurrent excitatory synapses), unspecific and stochastic, modeled as a homogeneous Poisson process with constant intensity *ν*_x_. Spike-triggered synaptic conductances are modeled as exponential functions, with fixed and equal conduction delays for all synapse types. The equations of the model dynamics, along with the numerical values for all parameters are summarized in S1 Table and S2 Table.

Following [33], the peak conductances were chosen such that the populations operate in a balanced, low-rate asynchronous irregular regime when driven solely by background input. For this purpose, we set 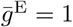 and 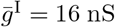, giving rise to average firing rates of ~3 Hz, CV_ISI_ ∈ [1.0, 1.5] and CC ≤ 0.01 in the first two modules of the networks, as described in the previous sections.

### Stimulus input and computational tasks

We evaluate the information processing capabilities of the different networks on simple linear and nonlinear computational tasks. For this purpose, the systems are presented with a sequence of stimuli {*S*_1_, *S*_2_,…} ∈ *S*, of finite total length *T* and comprising |*S*| different stimuli.

Each stimulus consists of a set of 800 Poisson processes at a fixed rate *ν*_stim_ = λ * *ν*_x_ and fixed duration of 200 ms, mimicking sparse input from an external population of size *N*^E^ (Fig 1C). These input neurons are mapped to randomly chosen, but stimulus-specific sub-populations of *ϵN*^E^ excitatory and *ϵN*^I^ inhibitory neurons in the first module *M*_0_, which we denote the *input module*. Unless otherwise stated, we set λ = 3, resulting in mean firing rates ranging between 2-8 spikes/s across the modules.

To sample the population responses for each stimulus in the sequence, we collect the responses of the excitatory population in each module *M_i_* at fixed time points *t*\*, relative to stimulus onset (with *t*\* = 200 ms, unless otherwise stated). These activity vectors are then gathered in a state matrix *X_M_i__* ∈ ℝ^*N*^E^×*T*^. In some cases, the measured responses are quantified using the low-pass filtered spike trains of the individual neurons, obtained by convolving them with an exponential kernel with *τ* = 20 ms and temporal resolution equal to the simulation resolution, 0. 1 ms. However, for most of the analyses, we consider the membrane potential *V*_m_ as the primary state variable, as it is parameter-free and constitutes a more natural choice [34, 35].

### Classification of stimulus identity

In the simplest task, the population responses are used to decode the identity of the input stimuli. The classification accuracy is determined by the capacity to linearly combine the input-driven population responses to approximate a target output [24]:

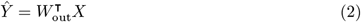

where *Ŷ* ∈ ℝ^*r*×*T*^ and *X* ∈ ℝ^*N*^E^×*T*^ are the collection of all readout outputs and corresponding states over all time steps *T* respectively, and *W*_out_ is the *N*^E^ × *r* matrix of output weights from the excitatory populations in each module to their dedicated readout units. We use 80% of the input data for training a set of *r* linear readouts to correctly classify the sequence of stimulus patterns in each module, where *r* = |*S*| is the number of different stimuli to be classified. Training is performed using ridge regression (*L*_2_ regularization), with the regularization parameter chosen by leave-one-out cross-validation on the training dataset. In the test phase, we obtain the predicted stimulus labels for the remaining 20% of the input sequence by applying the winner-takes-all (WTA) operation on the readout outputs *Ŷ*. Average classification performance is then measured as the fraction of correctly classified patterns.

### Nonlinear exclusive-or (XOR)

We also investigate the more complex XOR task, involving two parallel stimulus sources *S* and *S*′ injected into either the same or two separate input modules. Given stimulus sets *S* = {*S*_0_, *S*_1_} and 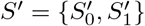, the task is to compute the XOR on the stimulus labels, i.e., the target output is 1 for input combinations 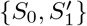 and 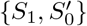, and 0 otherwise. In this case, computational performance is quantified using the point-biserial correlation coefficient (PBCC), which is suitable for determining the correlation between a binary and a continuous variable [33, 36, 37]. The coefficient is computed between the binary target variable and the analog (raw) readout output *Ŷ*(*t*), taking values in the [−1, 1] interval, with any significantly positive value reflecting a performance above chance.

### State separation analysis

To visualize and interpret the geometric arrangement of the population response vectors in the network’s state-space, we analyze the characteristics of a low-dimensional projection of the firing rate vectors obtained through principal component analysis (PCA), and evaluate how similar each data point (state vector projected onto the space spanned by the first 3 PCs) in one stimulus-specific cluster is to its own cluster compared to neighboring clusters. This is done by assigning a silhouette coefficient, ranging between [−1, 1], to each labeled sample of population activity during a single trial. A coefficient value close to 1 indicates that the data point is close to the mean of its assigned cluster (correct stimulus label), near 0 points to partially overlapping clusters, while negative values imply wrong cluster assignment.

### Numerical simulations and analysis

All numerical simulations were conducted using the Neural Microcircuit Simulation and Analysis Toolkit (NMSAT) v0.2 [4], a high-level Python framework for creating, simulating and evaluating complex, spiking neural microcircuits in a modular fashion. It builds on the PyNEST interface for NEST [38], which provides the core simulation engine. To ensure the reproduction of all the numerical experiments and figures presented in this study, and abide by the recommendations proposed in [39], we provide a complete code package that implements project-specific functionality within NMSAT (see S1 Appendix) using a modified version of NEST 2.12.0 [40].

## Results

Distributed information processing across multiple neural circuits requires, in a first instance, an accurate representation of the stimulus identity and a reliable propagation of this information throughout the different modules. In the following section, we assess these capabilities using a linear classification task in a sequential setup (illustrated in Fig 1), and analyze the characteristics of population responses in the different modules. Subsequently, we look at how different network setups handle information from two concurrent input streams by examining their ability to perform nonlinear transformations on the inputs.

### Sequential transmission of stimulus representation

In networks with fully random projections (Fig 1A), stimulus information can be accurately decoded up to a maximum depth of 3, i.e. the first three modules in the sequential setup contain sufficient information to classify (significantly beyond chance level) which of the ten stimuli had been presented to the input module (see Methods for details of the stimulus generation and classification assessment). Whereas the first two modules, *M*_0_ and *M*_1_, achieve maximum classification performance with virtually no variance across trials (Fig 2A, plain bars), the accuracy of ≈ 0.55 observed in *M*_2_ indicates that the stimulus representations have become degraded. These results suggest that while random connectivity between the modules allows the input signal to reach *M*_2_, the population responses at this depth are already insufficiently discernible to propagate further downstream, with *M*_3_ entirely unable to distinctly represent the different stimuli.

**Fig 2.**
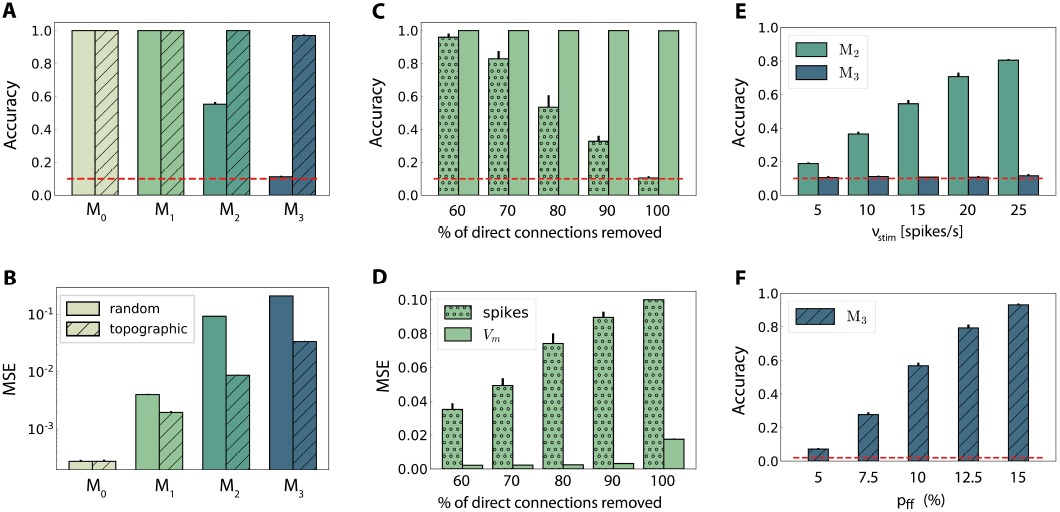
Stimulus classification in sequentially connected modular networks. **A, B**: Mean classification accuracy over |*S*| = 10 stimuli and corresponding mean squared error in each of the four modules in the random (plain bars) and topographic (hatched bars) conditions. **C, D**: Mean classification accuracy and corresponding mean squared error in *M*_1_ as a function of the number of direct projections (from neurons receiving direct stimulus input in *M*_0_ to neurons in *M*_1_) when decoding stimulus information from the low-pass filtered spike trains (stippled bars) and the membrane potential (plain bars). **E**: Classification accuracy in *M*_2_ and *M*_3_ decoded from the membrane potential as a function of the input intensity. **F**: Classification accuracy over |*S*| = 50 stimuli in *M*_3_ as a function of the connection density within the topographic projections. All panels show the mean and standard deviations obtained from ten simulations per condition.

Including structured projections in the system (Fig 1B) counteracts these effects, allowing stimulus information to be accurately transferred to the deeper modules (Fig 2A, hatched bars). This indicates that stimulus-specific topographic maps, whereby the neurons receiving direct stimulation at *M_i_* connect exclusively to another set of stimulus-specific neurons in the subsequent module (see Methods), play a critical role in the successful propagation of signals across multiple interacting sub-networks.

As computing the accuracy scores involves a nonlinear post-processing step (winner-takes-all, see Methods), we additionally verify whether this operation significantly biases the results by evaluating the mean squared error (MSE) between the raw readout outputs *Ŷ* and the binary targets *Y*. These MSE values, depicted in Fig 2B, are consistent: performance decays with depth for both network setups, with topography leading to significant computational benefits for all modules beyond the input module. In the following two sections, we investigate the factors influencing stimulus propagation and uncover the relationships between the underlying population dynamics and the system’s task performance.

### Modulating stimulus propagation

Since random networks provide no clearly structured feed-forward pathways to facilitate signal propagation, it is unclear how stimulus information can be read out as far as *M*_2_ (Fig 2A), considering the nonlinear transformations at each processing stage. However, by construction, some neurons in *M*_0_ that receive input stimulus directly also project (randomly) to *M*_1_. To assess the importance of these directed projections for information transmission, we gradually remove them and measure the impact on the performance in *M*_1_ (Fig 2C, D). The system shows substantial robustness with respect to the loss of such direct feed-forward projections, as the onset of the decline in performance only occurs after removing half of the direct synapses. Furthermore, this decay is observed almost exclusively in the low-pass filtered responses, while the accuracy of state representations at the level of membrane potentials remains maximal. This suggests that the populations in the input module are not only able to create internal representations of the stimuli through their recurrent connections, but also transfer these to the next module in an suitable manner. The different results obtained when considering spiking activity and sub-threshold dynamics indicate that the functional impact of recurrence is much more evident in the population membrane potentials.

It is reasonable to assume that the transmission quality in the two networks, as presented above, is susceptible to variations in the input intensity. For random networks, one might expect that increasing the stimulus intensity would enable its decoding in all four modules. Although stronger input does improve the classification performance in *M*_2_ (Fig 2E), this improvement is not visible in the last module. When varying the input rates between 5 and 25 spk/s, the accuracy increases linearly with the stimulus intensity in *M*_2_. However, the signal does not propagate to the last module in a decipherable manner (results remain at chance level), regardless of the input rate and, surprisingly, regardless of the representational accuracy in *M*_2_.

Previous studies have shown that, when structured feed-forward connections are introduced, the spiking activity propagation generally depends on both the synaptic strength and connection density along the structures, with higher values increasing the transmission success [16, 21]. To evaluate this in our model without altering the synaptic parameters, we increase the task difficulty and test the ability of the last module, *M*_3_, to discriminate 50 different stimuli. The results, shown in Fig 2F, exhibit a significantly lower performance for the initial topographic density of (7.5%), from ≈ 1 for ten stimuli (Fig 2A) to ≈ 0.3. This drop can be likely attributed to overlapping projections between the modules, since more stimulus-specific pathways naturally lead to more overlap between these regions, causing less discriminable responses. However, this seems to be compensated for by increasing the projection density, with stronger connectivity significantly improving the performance. Thus, our simulations corroborate these previous experiments: increasing the connection density within topographic maps increases the network’s computational capacity.

### Population activity and state separability

To ensure a perfect linear decoding of the input, population responses elicited by different stimuli must flow along well segregated, stimulus-specific regions in the network’s state-space (separation property, see [23]). In this section, we evaluate the quality of these input-state mappings as the representations are transferred from module to module, and identify population activity features that influence the networks’ computational capabilities in various scenarios.

When a random network is driven only by background noise, the activity in the first two modules is asynchronous and irregular, but evolves into a more synchronous regime in *M*_2_ (see example activity in Fig 3A left, and noise condition in Fig 3B). In the last module, the system enters a synchronous regime, which has been previously shown to negatively impact information processing by increasing redundancy in the population activity [33]. This excessive synchronization explains the increased firing rates, reaching ≈ 10 spk/s in *M*_3_ (Fig 3D). Previous works have shown that even weak correlations within an input population can induce correlations and fast oscillations in the network [31]. This phenomenon arises in networks with sequentially connected populations, and is primarily a consequence of an increase in shared pre-synaptic inputs between successive populations [15, 19, 42]. As the feed-forward projections gradually increase the convergence of the inter-module connections, the corresponding magnitude of post-synaptic responses also increase towards the deeper modules. Effectively stronger synapses then shift the network’s operating point away from the desired Poissonian statistics. This effect accumulates from module to module and gradually skews the population activity towards states of increased synchrony.

**Fig 3.**
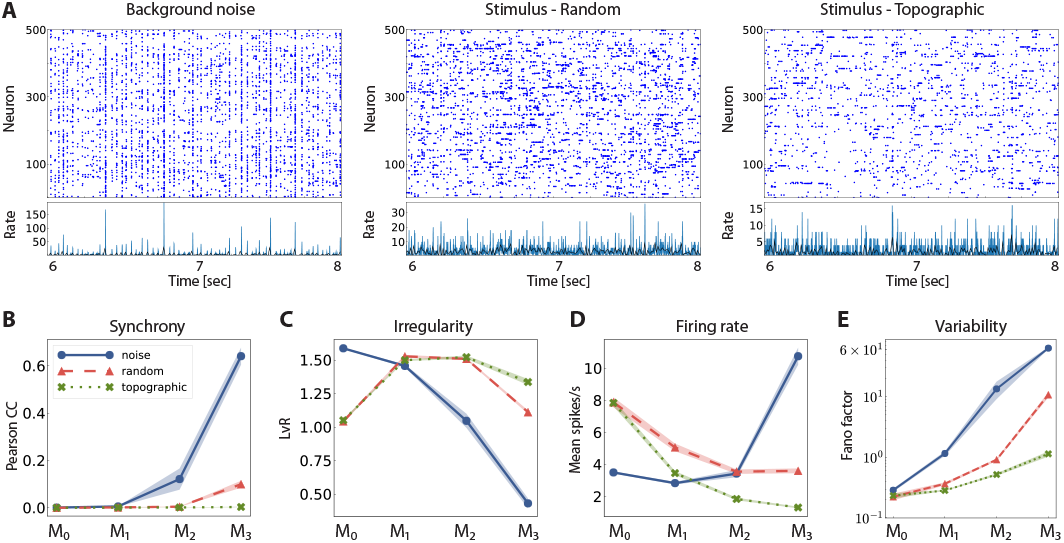
Network activity in three different scenarios: purely noise-driven (no stimulus); stimulus-driven with random feed-forward connections and stimulus-driven with structured, topographic projections. Top row (**A**) shows 2 seconds of spiking activity and the corresponding firing rates of 500 randomly chosen excitatory neurons in *M*_2_. The corresponding statistics across the excitatory populations are shown in the bottom row (**B-E**), for every module: **B** - synchrony (Pearson’s correlation coefficient, computed pairwise over spikes binned into 2 ms bins and averaged across 500 pairs); **C** - irregularity (measured as the revised local variation, LvR [41]); **D** - mean firing rate across the excitatory populations; and **E** response variability as measured by the Fano factor (FF) on the population-averaged firing rates (bin width 10 ms). All depicted statistics were averaged over ten simulations, each lasting 10 seconds, with ten input stimuli.

Compared to baseline activity, the presence of a patterned stimulus increases the irregularity in all modules except the very first one. This is visualized in the example activity plots in Fig 3A (center and right). Furthermore, active input substantially reduces the synchrony in the last two modules, allowing the system to globally maintain the asynchronous irregular regime (see random and topographic conditions in Fig 3B, C). Such alterations in the population response statistics during active processing have also been confirmed experimentally: *in vivo* recordings show that neuronal activity in awake, behaving animals is characterized by weak correlations and low firing rates in the presence of external stimuli [43, 44].

Despite the beneficial influence of targeted stimulation, it appears that random projections are not sufficient to entirely overcome the effects of shared input and excessive synchronization in the deeper modules (e.g. in *M*_3_, CC ≈ 0.12, with a correspondingly high firing rate). The existence of structured connectivity, through conserved topographic maps, on the other hand, allows the system to retain an asynchronous firing profile throughout the network. Whereas the more synchronous activity in random networks, coupled with a larger variability in the population responses (Fig 3E), contributes to their inability to represent the input in the deeper modules, topographic projections lead to more stable and reliable neuronal responses that enable the maintenance of distinguishable stimulus mappings, in line with the performance results observed in Fig 2A.

Furthermore, networks with structured connectivity are also more resource-efficient, achieving better performance with lower overall activity (Fig 3D). This can be explained by the fact that neurons receiving direct stimulus input in *M*_0_, firing at higher rates, project only to a restricted sub-population in the subsequent modules, thereby having a smaller impact on the average population activity downstream.

The above observations are also reflected in the geometric arrangement of the population response vectors, as visualised by the silhouette coefficients of a low-dimensional projection of their firing rates in Fig 4A (see Methods). As stimulus responses become less distinguishable with network depth, the coefficients decrease, indicating more overlapping representations. This demonstrates a reduction in the compactness of stimulus-dependent state vector clusters, which, although not uniformly reflected for all stimuli, is consistent across modules (only *M*_1_ and *M*_2_ shown). However, these coefficients are computed using only the first three principal components (PCs) of the firing rate vectors and are trial-specific. We can obtain a more representative result by repeating the analysis over multiple trials and taking into account the first ten PCs (Fig 4B). The silhouette scores computed in this way reveal a clear disparity between random and topographic network for the spatial segregation of the clusters, beginning with *M*_1_, in accordance with the classification performances (Fig 2A).

**Fig 4.**
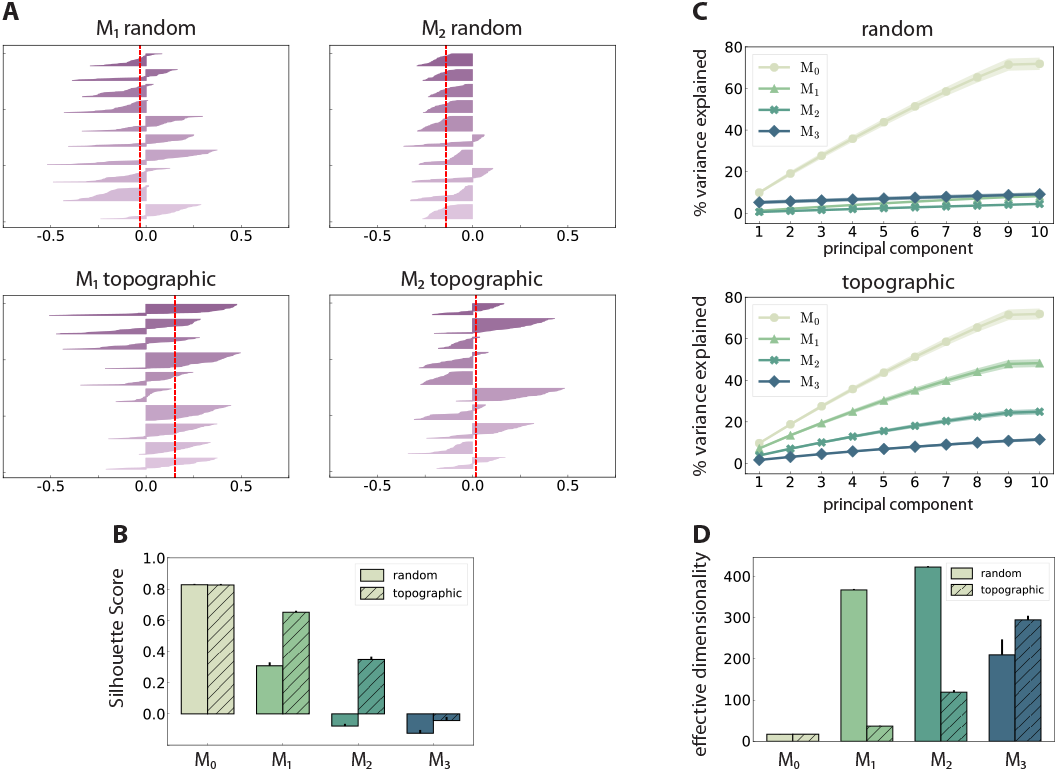
Spatial arrangement and cluster analysis of stimulus-specific state vectors. **A**: Distribution of silhouette coefficients for the stimulus-specific clusters in modules *M*_1_ and *M*_2_, computed in the space spanned by the first three principal components (PCs) of the firing rate vectors, and color-coded for the ten different stimuli used. The coefficients are sorted in descending order for each stimulus. The vertical lines in red represent the mean over all coefficients (silhouette score) in a single trial. **B**: Trial-averaged silhouette score calculated using the first ten PCs. **C**: Cumulative variance explained by the first ten PCs for random (top) and topographic (bottom) projections. **D**: Effective dimensionality of the state matrix computed on the firing rates (bin size 200 ms). All results are averaged over ten trials, each lasting 100 seconds (500 samples).

We can further assess the effectiveness with which the networks utilize their high-dimensional state-space by evaluating how many PCs are required to capture the majority of the variance in the data (Fig 4C). In the input module, where the stimulus impact is strongest, the variance captured by each subsequent PC is fairly constant (≈ 10%), reaching around 75% by the ninth PC. This indicates that population activity can represent the input in a very low-dimensional sub-space through narrow, stimulus-specific trajectories. In random networks, however, this trend is not reflected in the subsequent modules, where the first ten PCs account for less than 10% of the total variance.

There is thus a significant increase in the effective dimensionality [45] in the deeper modules (Fig 4D), a pattern which is also exhibited, to a lesser extent, in the topographic case. As the population activity becomes less entrained by the input, the deeper modules explore a larger region of the state-space. Whereas this tendency is consistent and more gradual for topographic networks, it is considerably faster in networks with unstructured projections, suggesting a quicker dispersion of the stimulus representations. Since in these networks the stimulus does not effectively reach the last module (Fig 2A), there is no de-correlation of the responses, and the elevated synchrony (Fig 3B) leads to a reduced effective dimensionality.

Overall, these results demonstrate that patterned stimuli push the population activity towards an asynchronous-irregular regime across the network, but purely random systems cannot sustain this state in the deeper modules. Networks with structured connectivity, on the other hand, display a more stable activity profile throughout the system, allowing the stimuli to propagate more efficiently and more accurately to all modules. Accordingly, the state representations are more compact and distinguishable, and these representations decay significantly slower with module depth than in random networks, in line with the observed classification results (Fig 2A).

### Memory capacity and stimulus sensitivity

As demonstrated above, both random and topographic networks are able to create unique representations of single stimuli in their internal dynamics and transfer these across multiple recurrent modules. In order to better understand the nature of distributed processing in these systems, it is critical to investigate how they retain information over time and whether representations of multiple, sequentially presented, stimuli can coexist in a superimposed manner, a property exhibited by cortical circuits as demonstrated by in vivo recordings [46].

To quantify these properties, we use the classification accuracy to evaluate how, for consecutive stimuli, the first stimulus decays and the second stimulus builds up (Fig 5A,B). For a given network configuration, the degree of overlap between the two curves indicates how long the system is able to retain useful information about both the previous and the present stimuli (Fig 5C). This analysis allows us to measure three important properties of the system: how long stimulus information is retained in each sub-network through reverberations of the current state; how long the network requires to accumulate sufficient evidence to classify the present input; and what are the potential interference effects between multiple stimuli.

**Fig 5.**
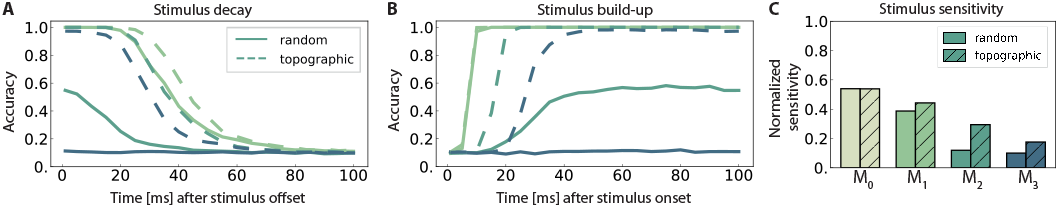
Stimulus sensitivity and temporal evolution of state representations, as indicated by the classification accuracy. **A** and **B** show the time course of the readout accuracy for the preceding and the current stimulus respectively, with *t* = 0 representing the offset of the previous and onset of the new stimulus. Curves depict the mean accuracy score over 5 trials, with linear interpolation of sampling offsets *t*_samp_ = 1, 5, 10, 15,…, 100 ms. Solid and dashed curves represent networks with random connectivity and topography respectively, color-coding according to modules (key in **C**). The stimulus sensitivity (**C**) is defined as the area below the intersection of corresponding curves from **A** and **B**, normalized with respect to maximum performance.

The decay in performance measured at increasing delays after stimulus offset (Fig 5A) shows how input representations gradually disappear over time (the *fading memory* property [14]). For computational reasons, only the first 100 ms are plotted, but the decreasing trend in the accuracy continues and invariably reaches chance level within the first 150 ms. This demonstrates that the networks have a rather short memory capacity which is unable to span multiple input elements, and that the ability to memorize stimulus information decays with network depth. Adding to the functional benefits of topographic maps, the memory curves reflect the higher overall accuracy achieved by these networks.

We further observe that the networks require exposure time to acquire discernible stimulus representations (Fig 5B). The time for classification accuracy to reach its maximum increases with depth, resulting in an unsurprising cumulative delay. Notably, topography enables a faster information build-up beginning with *M*_2_.

To determine the *stimulus sensitivity* of a population, we consider the extent of time where useful non-interfering representations are retained in each sub-network. This can be calculated as the area below the intersection of its memory and build-up curves. Following a similar trend to performance and memory, sensitivity to stimulus decreases with network depth and the existence of structured propagation pathways leads to clear benefits, particularly pronounced in the deeper modules (Fig 5C).

Overall, modules located deeper in the network forget faster and take longer (than the inter-module delays) to build up stimulus representations. No population is able to represent two sequential stimuli accurately for a significant amount of time (longer than 100 ms), although topographic maps improve memory capacity and stimulus sensitivity.

### Integrating multiple input streams

The previous section focuses on a single input stream, injected into a network with sequentially connected modules. Here, we examine the microcircuit’s capability to integrate information from two different input streams, in two different scenarios with respect to the location of the integration. The set-up and results are illustrated in Fig 6.

**Fig 6.**
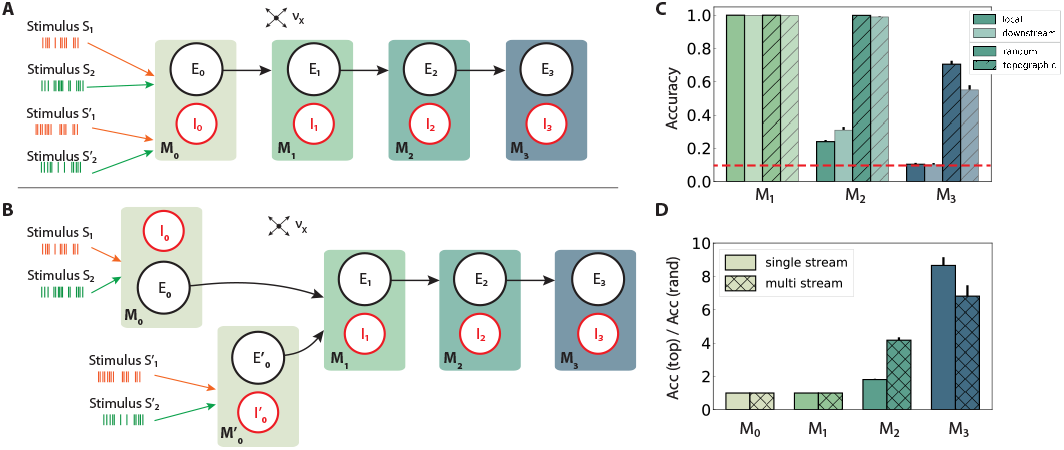
Schematic overview of information integration from two input streams (*S* and *S*′), and their performance on the stimulus classification task. **A, local integration:** sequential four-module set-up with two input streams injected into the first module, *M*_0_, where they are combined and transferred downstream. **B, downstream integration:** as in **A**, but with the first module divided into two separate sub-modules *M*_0_ and 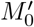, each receiving one stream as input and projecting to *M*_1_. Integration occurs in *M*_1_. Connection probabilities, weights and other parameters are identical to those in previous scenarios (see Fig 2A, B), with the exception of downstream integration (B): to keep the overall excitatory input to *M*_1_ consistent with local integration, projection densities to *M*_1_ from the input modules *M*_0_ and 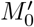 are scaled to *p*_ff_/2, while the remaining connections are left unchanged. **C**: classification accuracy of ten stimuli from one input stream, in modules *M*_1_ – *M*_3_. **D**: Relative performance gain in topographic networks, measured as the ratio of accuracy scores in the single and multiple stream (local integration) scenarios. Results are averaged over ten trials, with dark and light colors coding for local and downstream integration, respectively. The red dashed line represents chance level.

In a first step, the set-up from Fig 1A is extended with an additional input stream *S*′, without further alterations at population or connectivity level. The two stimulus sets, *S* and *S*′, are in principle identical, each containing the same number of unique stimuli and connected to specific sub-populations in the networks. Since the inputs are combined locally in the first module and the mixed information transferred downstream, we refer to this setup, visualized in Fig 6A, as *local integration*. In a second scenario (Fig 6B), each input stream is injected into a separate sub-module (*M*_0_ and 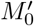), jointly forming the input module of the system. Here, computation on the combined input happens *downstream* from the first module, with the aim of simulating the integration of information that originated from more distant areas and had already been processed by two independent microcircuits.

Adding a second input stream significantly affects the network activity and the stimulus representations therein, which now must produce distinguishable responses for two stimuli concurrently. Compared to the same setup with a single input source (Fig 2A), the performance degrades in both random and topographic networks starting with *M*_2_ (Fig 6C, D). This suggests that the mixture of two stimuli results in less separable responses as the two representations interfere with each other, with structured connectivity again proving to be markedly beneficial. These benefits become clearer in the deeper modules, as demonstrated in Fig 6D where the effects of topography can lead to an 8-fold gain in task accuracy in *M*_3_.

As the spatio-temporal structure of the stimuli from both sources are essentially identical, it is to be expected that the mixed responses contain the same amount of information about both inputs. This is indeed the case, as reflected by comparable performance results when decoding from the second input stream (S1 Figure).

Interestingly, the location of the integration appears to play no major role for random networks. In networks with topographic maps, however, local integration improves the classification accuracy by around 25% in the last module compared to the downstream case. In the next section we investigate whether this phenomenon is set-up and task specific, or reflects a more generic computational principle.

### Local integration improves non-linear computation

In addition to the linear classification task discussed above, we analyze the ability of the circuit to extract and combine information from the two concurrent streams in a more complex, nonlinear fashion. For this, we trained the readouts on the commonly used non-linear XOR task described in the Methods section.

We observe that the networks’ computational capacity is considerably reduced compared to the simpler classification task, most noticeably in the deeper modules (Fig 7A). Although information about multiple stimuli from two input streams could be reasonably represented and transferred across the network, as shown in Fig 6C, it is substantially more difficult to perform complex transformations on even a small number of stimuli. This is best illustrated in the last module of topographic networks, where the stimulus identity can still be decoded with an accuracy of 70% (Fig 6C), but the XOR operation yields performance values close to chance level (PBCC of 0).

**Fig 7.**
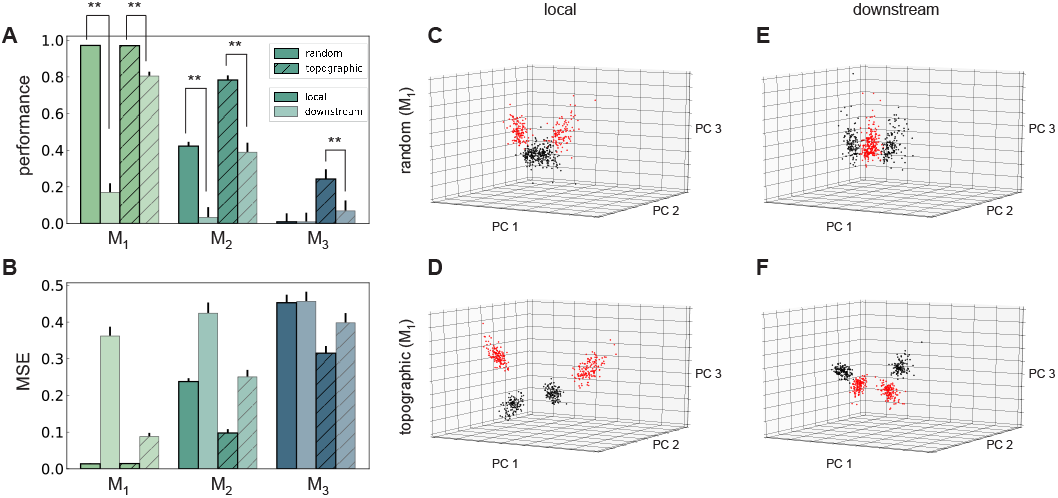
Performance and state-space partitioning in the XOR task. **(A)** Task performance on the XOR task measured using the point-biserial correlation coefficient (PBCC) between the XOR on the labels from the two input streams (target) and the raw readout values computed from the membrane potentials. **(B)** Corresponding mean squared error. Results are averaged over 10 trials, with 2000 training samples and 500 testing samples in each trial. **C-F:** PCA projections of 500 data points (low-pass filtered responses) in module *M*_1_, for the four combinations with respect to feed-forward connectivity and location of integration. Same axes in all panels. Colors code the target value, i.e., XOR on the stimulus labels from the two input streams. Random (left) and topographic (right) connectivity with local **(C,E)** and downstream **(D,F)** integration.

In contrast to the identity recognition, for XOR it is clearly more advantageous to fuse the two input streams in the first module (locally), rather than integrating only in *M*_1_ (Fig 7A,B). The differences in performance are statistically significant (two-sided Kolmogorov-Smirnov (KS) test 1.0, p-value < 0.01 for *M*_1_ and *M*_2_, and KS-test 0.9 with p-value < 0.01 for *M*_3_ in topographic networks) and consistent in every scenario and all modules from *M*_1_ onwards, with the exception of *M*_3_ in random networks.

One can gain a more intuitive understanding of the networks’ internal dynamics by looking at the state-space partitioning (Fig 7C-F), which reveals four discernible clusters corresponding to the four possible label combinations. These low-dimensional projections illustrate two key computational aspects: the narrower spread of the clusters in topographic networks (Fig 7D, F) is an indication of their greater representational precision, while the significance of the integration location is reflected in the collapse along the third PC in the downstream scenario (Fig 7F). To a lesser extent, these differences are also visible for random networks (Fig 7C, E). A more compact representation of the clustering quality using silhouette scores, consistent with these observations, is depicted in S2 Figure.

Altogether, these results suggest that it is computationally beneficial to perform non-linear transformations locally, as close to the input source as possible, and then propagate the result of the computation downstream instead of the other way around. The results were qualitatively similar for both the low-pass filtered spike trains and the membrane potential (see S3 Figure). To rule out any possible bias arising from re-scaling the feed-forward projections to *M*_1_ in the downstream scenario, we also ensured that these results still hold when each of the input sub-modules *M*_0_ and 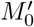 projected to *M*_1_ with the same unscaled probability *p*_ff_ as in Fig 1B (see S4 Figure).

### Effective dimensionality depends on the architecture of stimulus integration

Previous studies have suggested that non-linear integration of multiple input streams is associated with high response dimensionality compared to areas in which little or only linear interactions occur [47, 48]. To assess whether these predictions hold in our model, we consider different stimulus integration schemes and investigate whether the effective response dimensionality correlates with XOR accuracy, which is used to quantify the non-linear transformations performed by the system.

For simplicity, we focus only on random networks. To allow a better comparison between the integration schemes introduced in Fig 6, we explore two approaches to gradually interpolate the downstream scenario towards the local one in an attempt to approximate its properties. First, we distribute each input stream across the two segregated input sub-modules *M*_0_ and 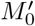, referred to as *mixed input* (Fig 8A). Second, we maintain the input stream separation but progressively merge the two sub-modules into a single larger one by redistributing the recurrent connections (Fig 8B). We call this scenario *mixed connectivity*.

**Fig 8.**
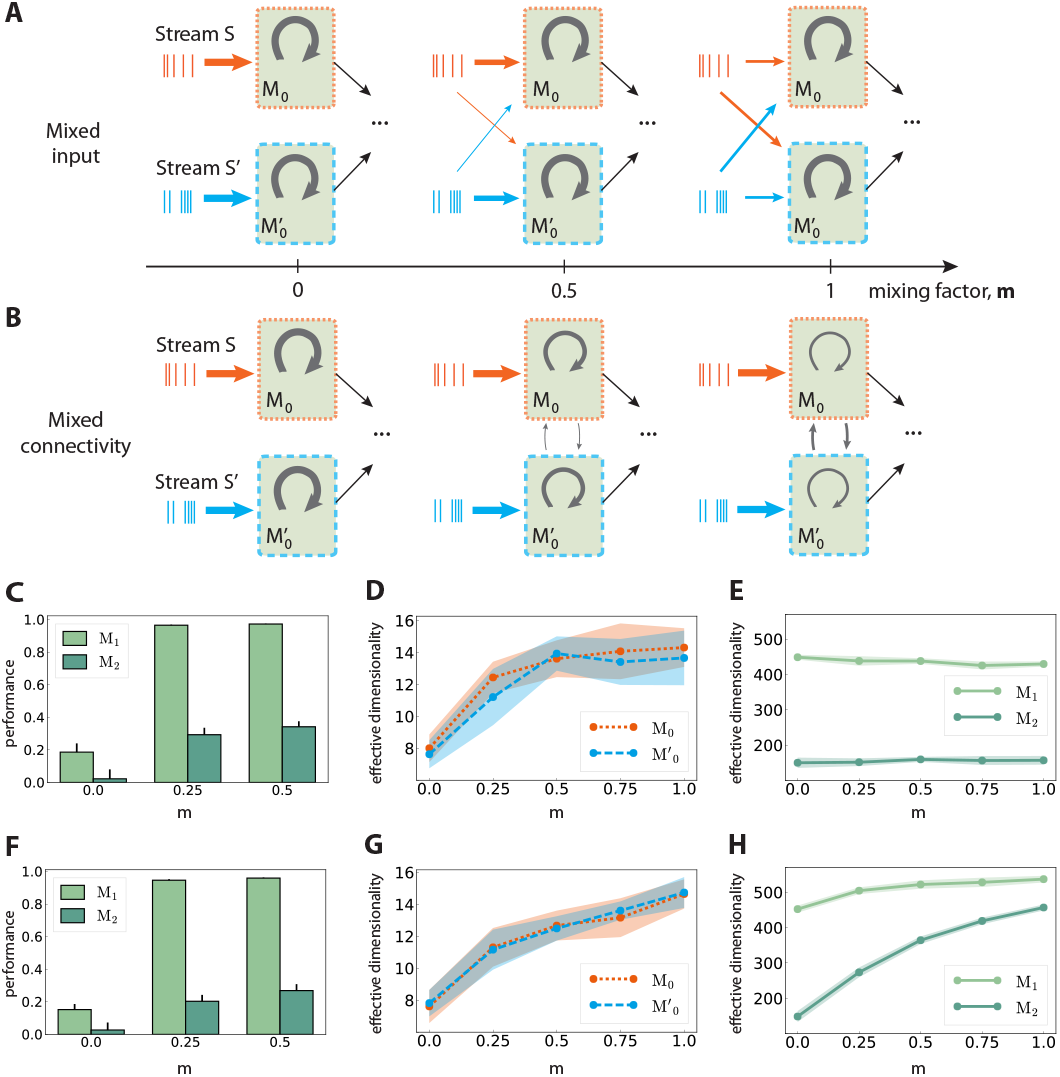
Mixed multi-stream integration and neural dimensionality in random networks. **A:** Downstream integration (only *M*_0_ shown) with progressive redistribution of the input across sub-modules *M*_0_ and 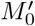. The mixing ratio is parametrized by *m* ∈ [0, 1], with *m* =0 representing two information sources mapped exclusively onto the corresponding sub-module, and *m* = 1 indicating equally distributed inputs across the sub-modules. The overall input to the network is kept constant. Arrow thickness indicates connection density. **B:** As in (**A**), but gradually connecting *M*_0_ and 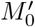 while keeping the input streams separated. Here, *m* controls the ratio of inter- and intra-module connection probabilities, with the total number of connections kept constant. **C:** XOR performance as a function of *m* for mixed input; **D:** Corresponding effective dimensionality in the input modules *M*_0_ and 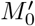; **E:** Corresponding effective dimensionality in the deeper modules *M*_1_ and *M*_2_. **F-H:** as in (**C-E**), but for mixed connectivity. Effective dimensionality is calculated for the membrane potential (mean and standard deviation over 10 trials, calculated using the first 500 PCs).

Relating these two scenarios is the *mixing factor* (*m*), which controls the input mapping or the connectivity between the sub-modules, respectively. A factor of 0 represents separated input sources and disconnected sub-modules as in Fig 6B; a value of *m* = 1 indicates that the input modules mix contributions from both sources equally (for mixed input), or that intra and inter-module connectivity for *M*_0_ and 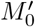 are identical (for mixed connectivity). In both cases, care was taken to keep the overall input to the network unchanged, as well as the average in- and out-degree of the neurons.

Combining information from both input streams already in the first sub-modules (*m* > 0), either via mixed input or mixed connectivity, significantly increases the task performance after convergence in the deeper modules. This is illustrated in Fig 8C, F, with *m* > 0.5 yielding similar values. Despite comparable gains in the nonlinear computational performance, the underlying mechanisms appear to differ in the two mixing approaches, as detailed in the following.

In *M*_0_ and 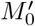, the effective dimensionality of the neural responses increases monotonically with the amount of information shared between the two modules (Fig 8D, G). This is expected, since the sub-modules are completely independent initially (*m* = 0) and can therefore use more compact state representations for single stimuli. However, diverging patterns emerge after convergence in *M*_1_. While the dimensionality does increase with the coefficient *m* in the mixed connectivity scenario (Fig 8H), it remains fairly constant in the mixed input case (Fig 8E), despite comparable task performance. Thus, complex non-linear transformations do not necessarily involve the exploration of larger regions of state-space, but can also be achieved through more efficient representations.

These results also demonstrate the difficulty in defining a clear relation between the ability of the system to perform nonlinear transformations on the input and its response dimensionality. Particularly in the case of larger networks involving transmission across multiple modules, the effective dimensionality can depend on the system’s architecture, such as the input mapping and connectivity structure in the initial stages.

## Discussion

Real-time interactions between a dynamic environment and a modular, hierarchical system like the mammalian neocortex strictly requires efficient and reliable mechanisms supporting the acquisition and propagation of adequate internal representations. Stable and reliable representations of relevant stimulus features must permeate the system in order to allow it to perform both local and distributed computations online. Throughout this study, we have analysed the characteristics of state representations in modular spiking networks and the architectural and dynamical constraints that influence the system’s ability to retain, transfer and integrate stimulus information.

We have considered models of local microcircuits as state-dependent processing reservoirs whose computations are performed by the systems’ high-dimensional transient dynamics [49, 50], acting as a temporal expansion operator, and investigated how the features of long-range connectivity in a modular architecture influence the system’s overall computational properties. By considering the network as a large modular reservoir, composed of multiple sub-systems, we have explored the role played by biologically-inspired connectivity features (conserved topographic projections) in the reliable information propagation across the modules, as well as the underlying dynamics that support the development and maintenance of such internal representations.

In addition to examining the temporal dynamics of the information transferred between sequentially connected modules, we have explored how different network characteristics enable information integration from two independent sources in a computationally useful manner. In these experiments, structural differences in the network were proven to greatly influence the dynamics and the downstream computation when combining inputs from two independent sources. In addition to the inter-module connectivity, the ability of the downstream modules to non-linearly combine the inputs was shown to depend on the location where the input converges, as well as on the extent to which the different input streams are mixed in the initial modules. We therefore anticipate that degree of mixed selectivity in early sensory stages is predictive of the computational outcome in deeper levels, particularly for non-linear processing tasks, as we describe in greater detail below.

### Representation transfer in sequential hierarchies

The proficiency of randomly coupled spiking networks (see e.g. [23, 33, 50]) demonstrates that random connectivity can be sufficient for local information processing. Successful signal propagation over multiple modules, however, appears to require some form of structured pathways for accurate and reliable transmission. Our results suggest that these requirements can be achieved by embedding simple topographic projections in the connectivity between the modules. Such mechanisms might be employed across the brain for fast and robust communication, particularly (but not exclusively) in the early sensory systems, where real-time computation is crucial and where the existence of topographic maps is well supported by anatomical studies [26, 51].

Purely random feed-forward connectivity allowed stimulus information to be decoded only up to the third module, whereas incorporating topographic projections ensured almost perfect accuracy in all modules (Fig 2A, B). These differences could be attributed to a decrease in the specificity of stimulus tuning with network depth, which is much more prominent for random networks (Fig 4). This result suggests that accurate information transmission over longer distances is not possible without topographic precision, thus uncovering an important functional role of this common anatomical feature.

Moreover, topography was shown to counteract the shared-input effect which leads to the development of synchronous regimes in the deeper modules. By doing so, stimulus information is allowed to propagate not only more robustly, but also more efficiently with respect to resources, in that the average spike emission is much lower (Fig 3D, E). Nevertheless, as the stimulus intensity invariably fades with network depth, the deeper modules capture fewer spatio-temporal features of the input and their response dimensionality increases. This process is clearer in random networks (Fig 4C, D), a further indication that topography enforces more stereotypical, lower-dimensional and stimulus-specific response trajectories. The input-state mappings are also retained longer and built up more rapidly in topographic networks (Fig 5).

### Network architecture and input integration

In biological microcircuits, local connections are complemented by long range projections which either stem from other cortical regions (cortico-cortical), or from different sub-cortical nuclei (e.g., thalamo-cortical). These different projections carry different information content and thus require the processing circuits to integrate multiple input streams during online processing. The ability of local modules to process information from multiple sources simultaneously and effectively is thus a fundamental building block of cortical processing.

Including a second input source into our sequential networks leads to less discriminable responses, as reflected in a decreased classification performance (Fig 6C). Integrating information from the two sources as early as possible in the system (i.e. in the modules closest to the input) was found to be clearly more advantageous for non-linear computations (Fig 6C) and, to a lesser extent, also to linear computations. For both tasks, however, topographic networks achieved better overall performance.

One of our main results thus suggests that computing locally, within a module, and transmitting the outcome of such computation (local integration scenario) is more effective than transmitting partial information and computing downstream. Accordingly, even a single step of non-linear transformation on individual inputs (downstream integration scenario) hinders the ability of subsequent modules to exploit non-trivial dependencies and features in the data. Therefore, it might be more efficient to integrate information and extract relevant features within local microcircuits that can act as individual computational units (e.g. cortical columns [5]). By combining the inputs locally, the population can create more stable representations which can then be robustly transferred across multiple modules. We speculate that in hierarchical cortical microcircuits, contextual information (simply modeled as a second input stream here) must be present in the early processing stages to enable more accurate computations in the deeper modules. This could, in part, explain the role of feedback connections from higher to lower processing centers.

### Degree of mixed selectivity predicts computational performance

We have further shown that the effective dimensionality of the neural responses does not correlate with the non-linear computational capabilities, except in the very first modules (Fig 8). These insights are in agreement with previous studies [47, 48], based on fMRI data that predict a high response dimensionality in areas involved in nonlinear multi-stream integration, and lower in areas where inputs from independent sources do not interact at all or solely overlap linearly. These studies considered single circuits driven by input from two independent sources, focusing on the role of mixed selectivity neurons in the convergent population. Mixed selectivity refers to neurons being tuned to mixtures of multiple task-related aspects [52, 53], which we approximated as a differential driving of the neurons with a variable degree of input from both sources.

Although we did not specifically examine mixed selectivity at a single neuron level, one can consider both the mixed input and mixed connectivity scenarios (Fig 8A and B, respectively) to approximate this behavior at a population level. This is particularly the case for the input sub-modules *M*_0_ and 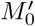, where the network’s response dimensionality, as expected, increases with the mixing ratio (Fig 8D, G). However, the different results we obtained for the deeper modules (Fig 8E, H), suggest that the effective dimensionality measured at the neuronal level is not a reliable evidence for non-linear processing in downstream convergence areas (despite similar performance), but instead depends on how information is mixed in the early stages of the system. Further research in this direction, possibly resorting to multimodal imaging data, is needed to determine a clear relation between functional performance, integration schemes and response dimensionality.

In our models, the task performance improved (and plateaued) with increased mixing factors, suggesting no obvious computational disadvantages for large factor values. While this holds for the discrimination capability of the networks, we did not address their ability to generalize. Since the sparsity of mixed selectivity neurons has been previously shown to control the discrimination-generalization trade-off, along with the existence of an optimal sparsity for neural representations [48], it would be interesting to analyze the effect of this parameter more thoroughly in the context of hierarchical processing.

Based on the presented findings, we expect that the degree of mixed selectivity in early sensory stages can predict the computational performance in the deeper levels, particularly for non-linear processing tasks. This might be the case for some components in the initial stages of visual processing, for instance when multiple features are combined. Whereas the retinotopic maps are mostly conserved in the primary visual cortex [54, 55], these gradually overlap (approximated in the mixed input scenario) in the subsequent areas, giving rise to more complex receptive fields and tuning properties [56]. Our results suggest that topographic maps may play a vital role in balancing between accurate transmission of state representations as well as controlling where and how information is integrated.

Despite the limitations of our models, we have highlighted the importance of biologically plausible structural patterning for information processing in modular spiking networks. Even simple forms of topography were shown to significantly enhance computational performance in the deeper modules. Additionally, architectural constraints have a considerable impact on the effectiveness with which different inputs are integrated, with early mixing being clearly advantageous and highlighting a possibly relevant feature of hierarchical processing. Taken together, these results provide useful constraints for building modular systems composed of spiking balanced networks that enable accurate information transmission.

### Limitations and future work

Our analysis consisted of a relatively simple implementation both in terms of the microcircuit composition and the characteristics of topographic maps. Even though abstractions are required in any modelling study, it is important to highlight the inherent limitations and drawbacks.

We have employed a simple process to embed topographic maps in unstructured networks (see Methods), whereby the map size (i.e. size of a population involved in a specific pathway) was kept constant in all modules. Cortical maps, however, exhibit more structured and complex spatial organization [51], characterized by a decrease in topographic specificity with hierarchical depth. This, in turn, is likely a consequence of increasingly overlapping projections and increasing map sizes and is considered to have significant functional implications (see e.g. [52]), which we did not explore in this study.

In addition, cortical systems also display an abundance of feedback loops that exhibit, similarly to the feed-forward cortico-cortical connections, a high degree of specificity and spatial segregation [3, 57]. Although their functional role is not entirely unambiguous and depends on specific functional interpretations, a recent study [20] found that these feedback projections have a destabilizing effect on long-range signal propagation. Failure to account for feedback projections will therefore limit the scope and generalizability of our models. Nevertheless, this limitation does not invalidate the main conclusions pertaining to the importance of structured projections in signal propagation and integration.

Ultimately, understanding the core principles of cortical computation requires bridging neuro-anatomy and physiology with cognitively relevant computations. The classification and XOR problems we have employed here provided a convenient method to investigate information transfer across multiple spiking modules, and allowed us to shed light on the functional implications of the wiring architecture. However, it is imperative that future works tackle more complex, behaviorally relevant tasks and possibly more detailed anatomical and physiological observations to help disentangle the nature of cognitive processing across cortical hierarchies.

## Acknowledgments

The authors gratefully acknowledge the computing time granted by the JARA-HPC Vergabegremium on the supercomputer JURECA at Forschungszentrum Jülich. We acknowledge partial support by the Erasmus Mundus Joint Doctoral Program EuroSPIN, the German Ministry for Education and Research 1041 (Bundesministerium für Bildung und Forschung) BMBF Grant 01GQ0420 to BCCN Freiburg, the Initiative and Networking Fund of the Helmholtz Association and the Helmholtz Portfolio theme “Supercomputing and Modeling for the Human Brain”.

## Supporting information

**S1 Table Tabular description of network model after [58]**.

**S2 Table Summary of all the numerical values for the relevant parameters of the system**.

**S1 Figure.**
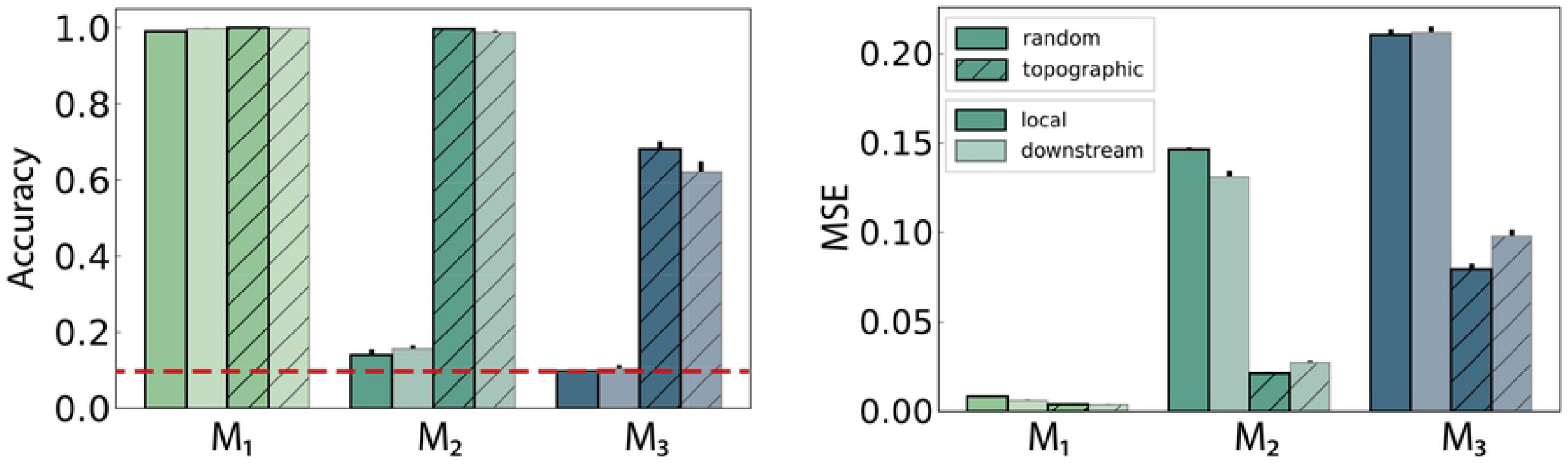
Figure Classification accuracy and MSE computed using ten stimuli from the second input stream, *S*′. There are no significant differences between local and downstream integration.

**S2 Figure.**
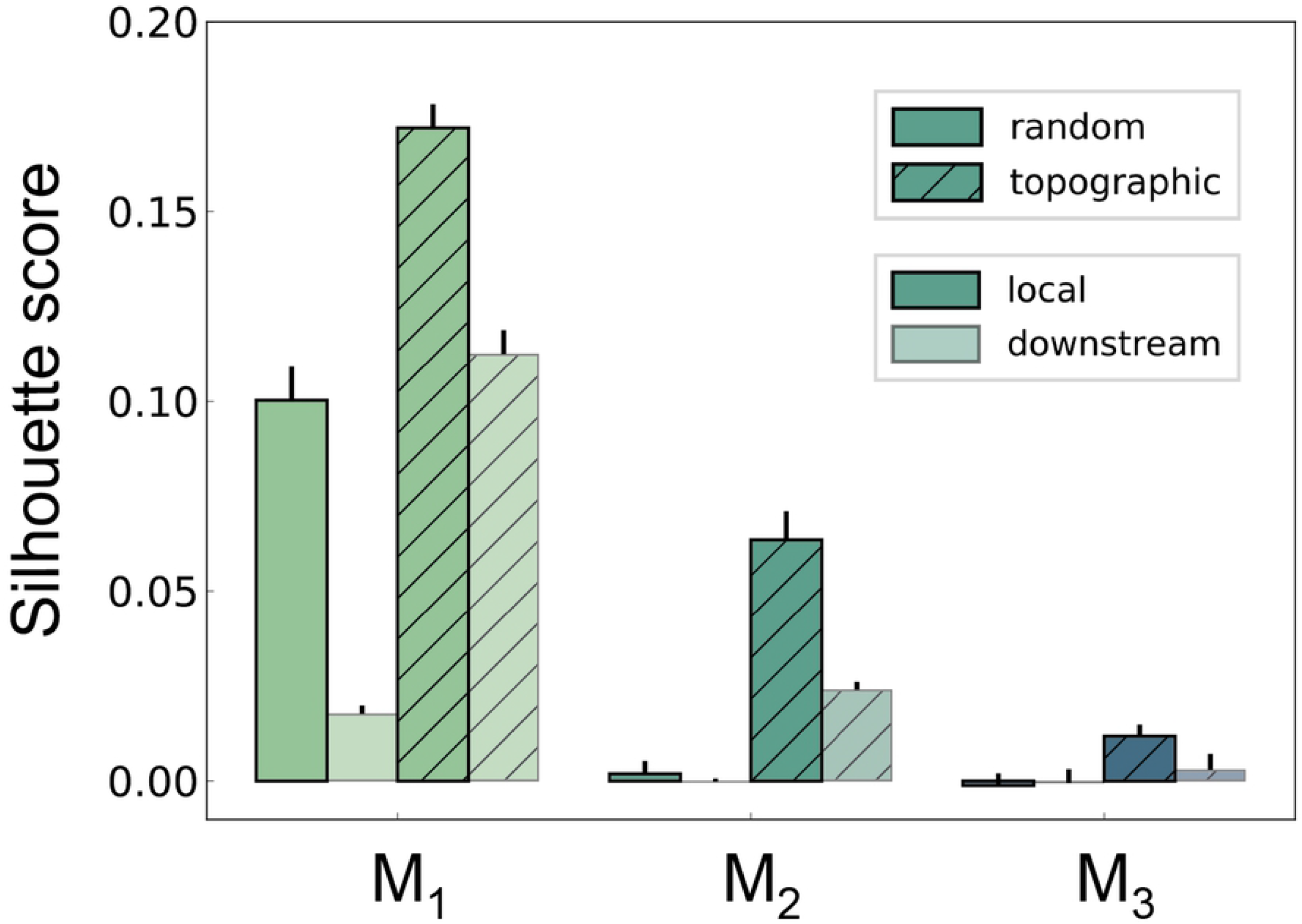
Silhouette score quantifying cluster separability in the XOR task. Scores are calculated in the space spanned by the first ten PCs, using the low-pass filtered spike trains.

**S3 Figure.**
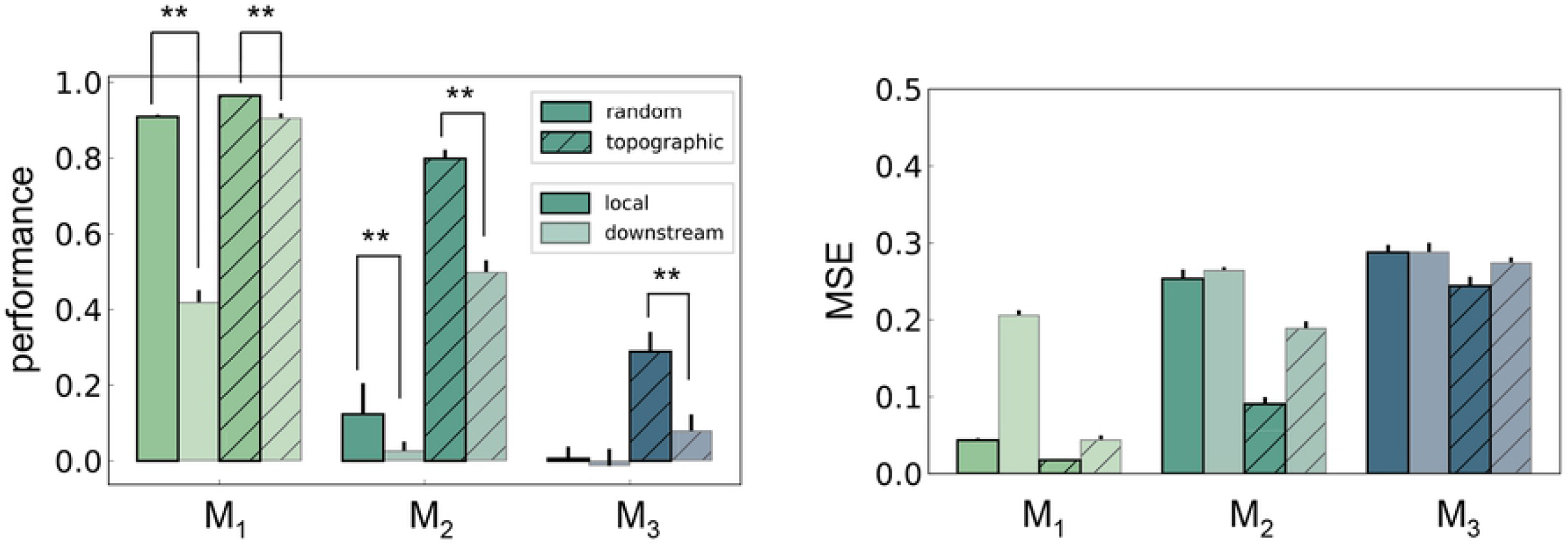
XOR performance computed on the low-pass filtered spike trains. The differences in performance are statistically significant, with local integration proving to be consistently more beneficial. These results are in agreement with the values computed using the membrane potentials as state variables.

**S4 Figure.**
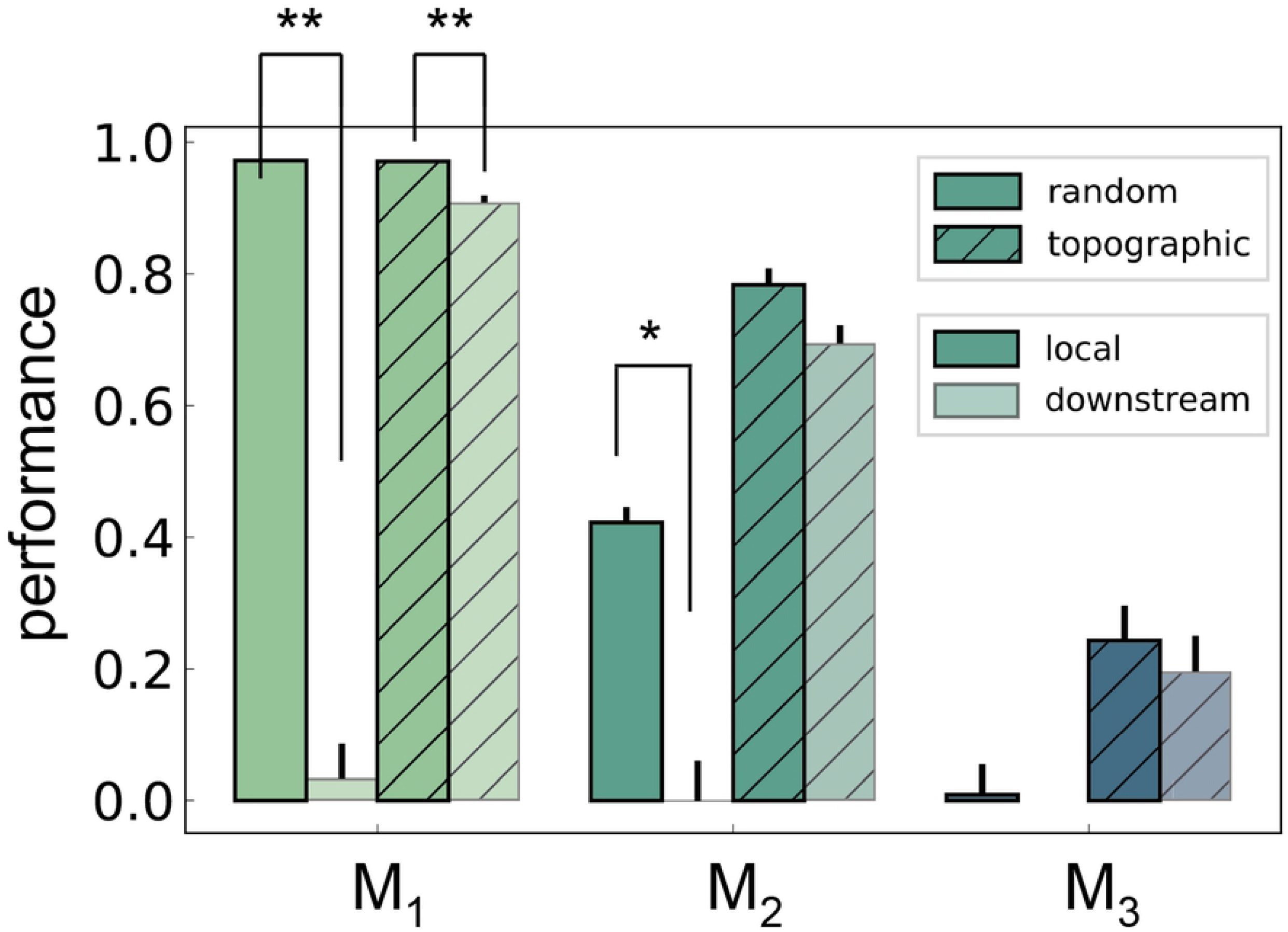
XOR performance for networks with non-scaled feed-forward projections between *M*_0_ → *M*_1_ and 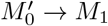 in the downstream integration scenario (Fig 6B). The denser connectivity (*p*_ff_ instead of *p*_ff_/2 as in Fig 7A) does not significantly alter the relative differences between local and downstream integration.

**S1 Appendix Reproducibility and replication**

**S1 Files Software package**.

## References

1. Felleman DJ, Van Essen DC. Distributed hierachical processing in the primate cerebral cortex. Cerebral Cortex. 1991;1(1):1–47. doi:10.1093/cercor/1.1.1.

2. Park HJ, Friston K. Structural and Functional Brain Networks: From Connections to Cognition. Science. 2013;342(6158):1238411–1238411. doi:10.1126/science.1238411.

3. Markov NT, Kennedy H. The importance of being hierarchical. Current Opinion in Neurobiology. 2013;23(2):187 – 194. doi:10.1016/j.conb.2012.12.008.

4. Duarte R, Seeholzer A, Zilles K, Morrison A. Synaptic patterning and the timescales of cortical dynamics. Current Opinion in Neurobiology. 2017;43:156–165. doi:10.1016/j.conb.2017.02.007.

5. Mountcastle VB. The columnar organization of the neocortex. Brain. 1997;120(4):701–722. doi:10.1093/brain/120.4.701.

6. Mountcastle V. An organizing principle for cerebral function: the unit model and the distributed system; 1978.

7. Duarte R. Expansion and State-Dependent Variability along Sensory Processing Streams. The Journal of Neuroscience. 2015;35(19):7315–7316. doi:10.1523/JNEUROSCI.0874-15.2015.

8. Macaluso E, Frith CD, Driver J. Modulation of Human Visual Cortex by Crossmodal Spatial Attention. Science. 2000;289(5482):1206–1208. doi:10.1126/science.289.5482.1206.

9. Keller G, Bonhoeffer T, Hübener M. Sensorimotor Mismatch Signals in Primary Visual Cortex of the Behaving Mouse. Neuron. 2012;74(5):809 – 815. doi:https://doi.org/10.1016/j.neuron.2012.03.040.

10. Shuler MG, Bear MF. Reward Timing in the Primary Visual Cortex. Science. 2006;311(5767):1606–1609. doi:10.1126/science.1123513.

11. Murray JD, Bernacchia A, Freedman DJ, Romo R, Wallis JD, Cai X, et al. A hierarchy of intrinsic timescales across primate cortex. Nature neuroscience. 2014;17(12):1661.

12. Miller KD. Canonical computations of cerebral cortex. Current Opinion in Neurobiology. 2016;37:75–84. doi:10.1016/j.conb.2016.01.008.

13. LeCun Y, Bengio Y, Hinton G. Deep learning. Nature. 2015;521(7553):436–444. doi:10.1038/nature14539.

14. Maass W, Natschläger T, Markram H. Fading memory and kernel properties of generic cortical microcircuit models. Journal of Physiology Paris. 2004;98(4-6 SPEC. ISS.):315–330. doi:10.1016/j.jphysparis.2005.09.020.

15. Kumar A, Rotter S, Aertsen A. Conditions for propagating synchronous spiking and asynchronous firing rates in a cortical network model. The Journal of neuroscience: the official journal of the Society for Neuroscience. 2008;28(20):5268–80. doi:10.1523/JNEUROSCI.2542-07.2008.

16. Kumar A, Rotter S, Aertsen A. Spiking activity propagation in neuronal networks: reconciling different perspectives on neural coding. Nature reviews neuroscience. 2010;11(9):615.

17. Diesmann M, Gewaltig MO, Aertsen A. Stable propagation of synchronous spiking in cortical neural networks. Nature. 1999;402(6761):529–533. doi:10.1038/990101.

18. van Rossum MCW, Turrigiano GG, Nelson SB. Fast propagation of firing rates through layered networks of noisy neurons. The Journal of neuroscience: the official journal of the Society for Neuroscience. 2002;22(5):1956–1966. doi:22/5/1956 [pii].

19. Shadlen MN, Newsome WT. The Variable Discharge of Cortical Neurons: Implications for Connectivity, Computation, and Information Coding. Journal of Neuroscience. 1998;18(10):3870–3896. doi:10.1523/JNEUROSCI.18-10-03870.1998.

20. Joglekar MR, Mejias JF, Yang GR, Wang XJ. Inter-areal balanced amplification enhances signal propagation in a large-scale circuit model of the primate cortex. Neuron. 2018;98(1):222–234.

21. Vogels TP, Abbott LF. Signal Propagation and Logic Gating in Networks of Integrate-and-Fire Neurons. The Journal of Neuroscience. 2005;25(46):10786–10795. doi:10.1523/JNEUROSCI.3508-05.2005.

22. Vogels TP, Abbott LF. Gating multiple signals through detailed balance of excitation and inhibition in spiking networks. Nature Neuroscience. 2009;12(4):483–491. doi:10.1038/nn.2276.

23. Maass W, Natschläger T, Markram H. Real-Time Computing Without Stable States: A New Framework for Neural Computation Based on Perturbations. Neural Computation. 2002;14(11):2531–2560. doi:10.1162/089976602760407955.

24. Lukoševičius M, Jaeger H. Reservoir computing approaches to recurrent neural network training. Computer Science Review. 2009;3(3):127–149. doi:10.1016/j.cosrev.2009.03.005.

25. Zajzon B, Duarte R, Morrison A. Transferring State Representations in Hierarchical Spiking Neural Networks. In: 2018 International Joint Conference on Neural Networks (IJCNN); 2018. p. 1–9.

26. Kaas JH. Topographic maps are fundamental to sensory processing. Brain research bulletin. 1997;44(2):107–112.

27. Silver MA, Ress D, Heeger DJ. Topographic maps of visual spatial attention in human parietal cortex. Journal of neurophysiology. 2005;94(2):1358–1371.

28. Harris KD, Shepherd GMG. The neocortical circuit: themes and variations. Nature Neuroscience. 2015;18(2):170–181. doi:10.1038/nn.3917.

29. Hagler DJ, Sereno MI. Spatial maps in frontal and prefrontal cortex. NeuroImage. 2006;29(2):567–577. doi:10.1016/j.neuroimage.2005.08.058.

30. Thivierge JP, Marcus GF. The topographic brain: from neural connectivity to cognition. Trends in Neurosciences. 2007;30(6):251–259. doi:10.1016/j.tins.2007.04.004.

31. Brunel N. Dynamics of networks of randomly connected excitatory and inhibitory spiking neurons. Journal of Physiology Paris. 2000;94(5-6):445–463. doi:10.1016/S0928-4257(00)01084-6.

32. Kremkow J, Aertsen A, Kumar A. Gating of Signal Propagation in Spiking Neural Networks by Balanced and Correlated Excitation and Inhibition. Journal of Neuroscience. 2010;30(47):15760–15768. doi:10.1523/JNEUROSCI.3874-10.2010.

33. Duarte RC, Morrison A. Dynamic stability of sequential stimulus representations in adapting neuronal networks. Front Comput Neurosci. 2014;8(October):124. doi:10.3389/fncom.2014.00124.

34. van den Broek D, Uhlmann M, Fitz H, Duarte R, Hagoort P, Petersson KM. The best spike filter kernel is a neuron; 2017.

35. Duarte R, Uhlmann M, den van Broek D, Fitz H, Petersson KM, Morrison A. Encoding symbolic sequences with spiking neural reservoirs. In: 2018 International Joint Conference on Neural Networks (IJCNN); 2018. p. 1–8.

36. Haeusler S, Maass W. A statistical analysis of information-processing properties of lamina-specific cortical microcircuit models. Cerebral Cortex. 2007;17(1):149–162. doi:10.1093/cercor/bhj132.

37. Klampfl S, Maass W. Emergence of dynamic memory traces in cortical microcircuit models through STDP. The Journal of neuroscience: the official journal of the Society for Neuroscience. 2013;33(28):11515–29. doi:10.1523/JNEUROSCI.5044-12.2013.

38. Gewaltig MO, Diesmann M. NEST (NEural Simulation Tool). Scholarpedia. 2007;2(4):1430.

39. Pauli R, Weidel P, Kunkel S, Morrison A. Reproducing Polychronization: A Guide to Maximizing the Reproducibility of Spiking Network Models. Frontiers in Neuroinformatics. 2018;12:46. doi:10.3389/fninf.2018.00046.

40. Kunkel S, Morrison A, Weidel P, Eppler JM, Sinha A, Schenck W, et al. NEST 2.12.0; 2017.

41. Shinomoto S, Kim H, Shimokawa T, Matsuno N, Funahashi S, Shima K, et al. Relating neuronal firing patterns to functional differentiation of cerebral cortex. PLoS Computational Biology. 2009;5(7). doi:10.1371/journal.pcbi.1000433.

42. Tetzlaff T, Buschermöhle M, Geisel T, Diesmann M. The spread of rate and correlation in stationary cortical networks. Neurocomputing. 2003;52:949–954.

43. Ecker AS, Berens P, Keliris GA, Bethge M, Logothetis NK, Tolias AS. Decorrelated neuronal firing in cortical microcircuits. Science (New York, NY). 2010;327(5965):584–7. doi:10.1126/science.1179867.

44. Vaadia E, Haalman I, Abeles M, Bergman H, Prut Y, Slovin H, et al. Dynamics of neuronal interactions in monkey cortex in relation to behavioural events. Nature. 1995;373(6514):515–518. doi:10.1038/373515a0.

45. Mazzucato L, Fontanini A, La Camera G. Stimuli reduce the dimensionality of cortical activity. Frontiers in Systems Neuroscience. 2016;10(11). doi:10.3389/fnsys.2016.00011.

46. Nikolic D, Hausler S, Singer W, Maass W. Distributed fading memory for stimulus properties in the primary visual cortex. PLoS Biol. 2009;7(12):e1000260. doi:10.1371/journal.pbio.1000260.

47. Rigotti M, Fusi S. Estimating the dimensionality of neural responses with fMRI Repetition Suppression. arXiv preprint arXiv:160503952. 2016;.

48. Barak O, Rigotti M, Fusi S. The Sparseness of Mixed Selectivity Neurons Controls the Generalization–Discrimination Trade-Off. Journal of Neuroscience. 2013;33(9):3844–3856. doi:10.1523/JNEUROSCI.2753-12.2013.

49. Mante V, Sussillo D, Shenoy KV, Newsome WT. Context-dependent computation by recurrent dynamics in prefrontal cortex. Nature. 2013;503(7474):78–84. doi:10.1038/nature12742.

50. Sussillo D. Neural circuits as computational dynamical systems. Current Opinion in Neurobiology. 2014;25:156–163. doi:10.1016/j.conb.2014.01.008.

51. Bednar JA, Wilson SP. Cortical Maps. The Neuroscientist. 2016;22(6):604–617. doi:10.1177/1073858415597645.

52. Rigotti M, Barak O, Warden MR, Wang XJ, Daw ND, Miller EK, et al. The importance of mixed selectivity in complex cognitive tasks. Nature. 2013;497(7451):585–590. doi:10.1038/nature12160.

53. Warden MR, Miller EK. Task-dependent changes in short-term memory in the prefrontal cortex. Journal of Neuroscience. 2010;30(47):15801–15810.

54. Girman SV, Sauvé Y, Lund RD. Receptive Field Properties of Single Neurons in Rat Primary Visual Cortex. Journal of Neurophysiology. 1999;82(1):301–311. doi:10.1152/jn.1999.82.1.301.

55. Adams DL, Horton JC. A Precise Retinotopic Map of Primate Striate Cortex Generated from the Representation of Angioscotomas. Journal of Neuroscience. 2003;23(9):3771–3789. doi:10.1523/JNEUROSCI.23-09-03771.2003.

56. Hubel DH. Eye, brain, and vision / David H. Hubel. Scientific American Library; 1988.

57. Markov NT, Vezoli J, Chameau P, Falchier A, Quilodran R, Huissoud C, et al. Anatomy of hierarchy: Feedforward and feedback pathways in macaque visual cortex. Journal of Comparative Neurology. 2014;522(1):225–259. doi:10.1002/cne.23458.

58. Nordlie E, Gewaltig MO, Plesser HE. Towards reproducible descriptions of neuronal network models. PLoS computational biology. 2009;5(8):e1000456.

